# Temporal dynamics of competition between statistical learning and episodic memory in intracranial recordings of human visual cortex

**DOI:** 10.1101/2022.03.14.484293

**Authors:** Brynn E. Sherman, Kathryn N. Graves, David M. Huberdeau, Imran H. Quraishi, Eyiyemisi C. Damisah, Nicholas B. Turk-Browne

## Abstract

The function of long-term memory is not just to reminisce about the past, but also to make predictions that help us behave appropriately and efficiently in the future. This predictive function of memory provides a new perspective on the classic question from memory research of why we remember some things but not others. If prediction is a key outcome of memory, then the extent to which an item generates a prediction signifies that this information already exists in memory and need not be encoded. We tested this principle using human intracranial EEG as a time-resolved method to quantify prediction in visual cortex during a statistical learning task and link the strength of these predictions to subsequent episodic memory behavior. Epilepsy patients of both sexes viewed rapid streams of scenes, some of which contained regularities that allowed the category of the next scene to be predicted. We verified that statistical learning occurred using neural frequency tagging and measured category prediction with multivariate pattern analysis. Although neural prediction was robust overall, this was driven entirely by items that were subsequently forgotten. Such interference provides a mechanism by which prediction can regulate memory formation to prioritize encoding of information that could help learn new predictive relationships.

**Significance Statement:** When faced with a new experience, we are rarely at a loss for what to do. Rather, because many aspects of the world are stable over time, we rely upon past experiences to generate expectations that guide behavior. Here we show that these expectations during a new experience come at the expense of memory for that experience. From intracranial recordings of visual cortex, we decoded what humans expected to see next in a series of photographs based on patterns of neural activity. Photographs that generated strong neural expectations were more likely to be forgotten in a later behavioral memory test. Prioritizing the storage of experiences that currently lead to weak expectations could help improve these expectations in future encounters.

## Introduction

Long-term memory has a limited capacity, and thus a major goal of psychology and neuroscience has been to identify factors that determine which memories to store. Well-known factors include attention (***Aly and Turk-Browne, 2017***), emotion (***Dolcos et al., 2017***), motivation (***Dickerson and Adcock, 2018***), stress (***Goldfarb, 2019***), and sleep (***Cowan et al., 2021***). Here we propose a new factor that constrains long-term memory formation.

Beyond reliving the past, a key function of memory is that it allows us to predict the future (***Schacter et al., 2012***). When faced with a new experience, we draw on related experiences from the past to know what is likely to happen when and where (***De Brigard, 2014; Biderman et al., 2020***). This knowledge is the result of statistical learning, which identifies patterns or regularities in the environment that repeat over time (***Sherman et al., 2020; Endress and Johnson, 2021***) and form the basis of predictions (***De Lange et al., 2018***). We hypothesize that the availability of these predictions during encoding affects whether a new memory is formed. Namely, if one of the main objectives of long-term memory is to enable prediction, in the service of adaptive behavior, experiences that already generate a prediction may not need to be encoded. In contrast, experiences that yield uncertainty about what will happen next may be more important to store because they can help learn over time what should have been expected. Note that this is distinct from whether an experience being encoded was itself expected or unexpected, which also affects subsequent memory (***Greve et al., 2017; Bein et al., 2021***); rather, we argue that the process of generating a prediction based on the experience impedes its encoding.

We term this ability of an experience to generate a prediction its *predictive value*. There is some suggestive evidence for predictive value as an encoding factor. In a statistical learning study with images presented in temporal pairs, subsequent memory for the first item in a pair was impaired relative to unpaired control items (***Sherman and Turk-Browne, 2020***). Because the first item in a pair was always followed by the second item, it could have enabled a prediction of the second item and thus had predictive value.

However, this prior study was not able to link the predictive value of an item during encoding to subsequent memory for that item for several reasons. One issue is that it was unclear whether the memory impairment for the first item originated at the time of encoding or emerged in later stages such as consolidation or retrieval. For example, the first item might have been encoded well, but when this item was probed in the later memory test, its association with the second item interfered with recognition. The behavioral experiments in the prior study were equivocal, as prediction was not measured during encoding. An fMRI experiment provided some evidence of prediction during encoding — the category of the second item could be decoded during the first — but the poor temporal resolution fMRI muddied this interpretation. The decoded neural signals were recorded during or after the second item and shifted backward in time based on assumptions about the hemodynamic lag. Methods with better temporal resolution could provide more precise linking between neural signals and experimental events, allowing for more direct measurement of predictions.

Another issue with the prior study is that it only examined the relationship between prediction and encoding across participants. Average fMRI evidence for the category of second items during first items was negatively associated with overall memory performance for first items. However, this could reflect a generic individual difference — that participants who make more predictions tend to have worse memory — rather than prediction having a mechanistic effect on encoding. According to the latter account, whether a participant remembers or forgets a given item should depend on whether that item triggered a prediction during its encoding. This requires testing for a relationship between prediction and encoding across items within participant.

The present study addresses these issues to better establish predictive value as an encoding factor. We combine intracranial EEG (iEEG) with multivariate pattern analysis, allowing us to measure neural predictions in a time-resolved manner and link them to subsequent behavioral memory across trials. Epilepsy patients viewed a rapid stream of scene photographs across blocks of a statistical learning task. The scenes consisted of unique exemplars from various categories (e.g., beaches, mountains, waterfalls) that differed by block. In the Random blocks, the order of “control” (condition X) categories from which the exemplars were drawn was random. In the Structured blocks, the categories were paired such that exemplars from “predictive” (condition A) categories were always followed by exemplars from “predictable” (condition B) categories (***Figure 1***A). Patients were not informed of these conditions or the existence of category pairs, which they learned incidentally through exposure (***Brady and Oliva, 2008***). The items from each category were presented in sub-blocks that changed after four presentations (***Figure 1***B). After both blocks, patients completed a recognition memory test for the exemplars from the stream.

**Figure 1.**
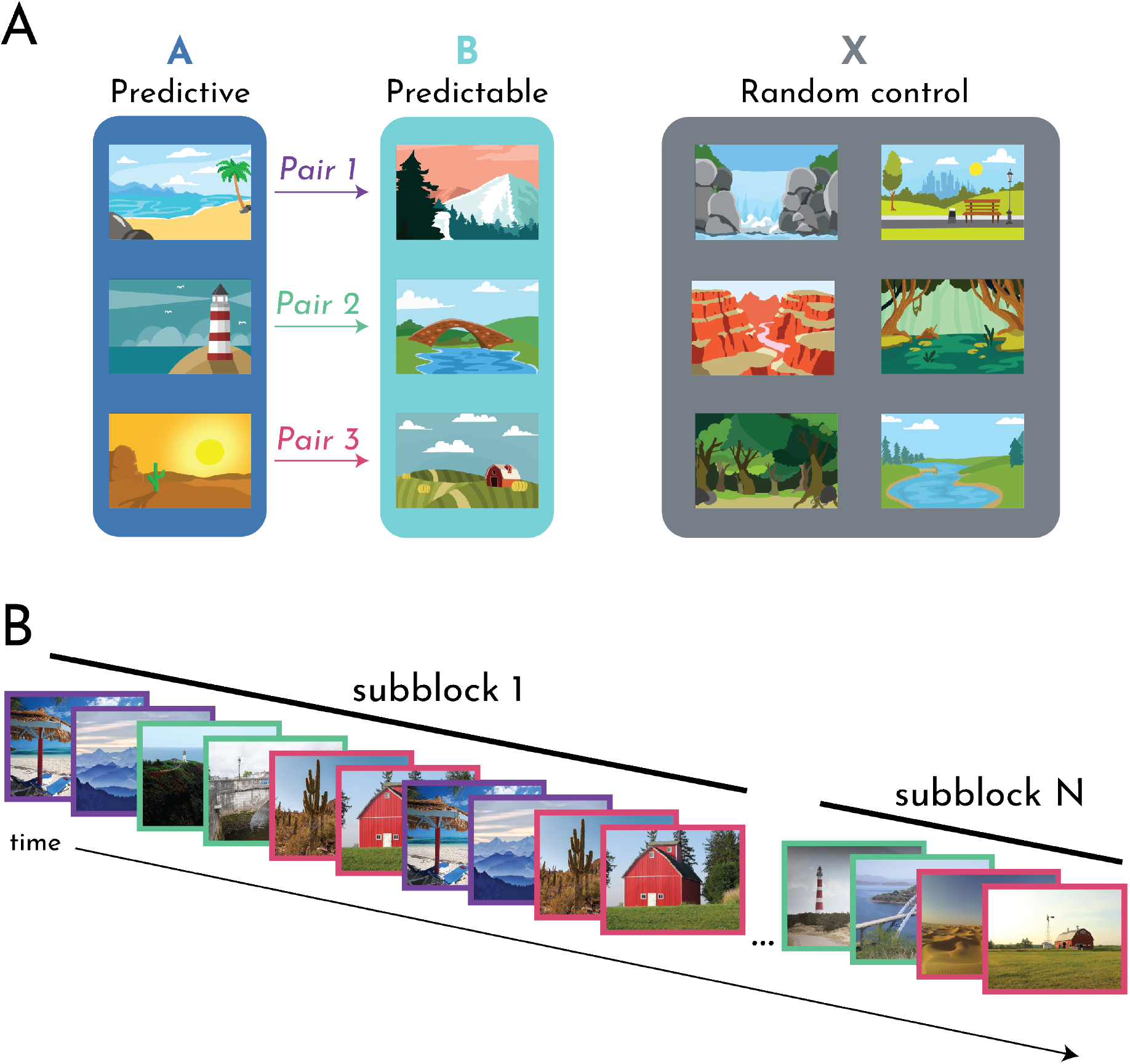
Task design. (A) Example scene category pairings for one participant. Three of 12 categories were assigned to condition A. Each A category was reliably followed by one of three other categories assigned to condition B to create pairs. The remaining six categories assigned to condition X were not paired. Participants viewed the A and B (Structured) and X (Random) categories in separate blocks of the task. (B) Example stimuli from the Structured block. Participants passively viewed a continuous stream of scenes. Each scene was shown for 267 ms, followed by an ISI of 267 ms with only a fixation cross on the screen. The stream was segmented into subblocks. The same exemplar of each category was presented four times per subblock, and new exemplars were introduced for the next subblock. For the Structured block, the category pairs remained consistent across subblocks. Category pairs are denoted by a colored frame, corresponding to the A-B pairs (and colored arrows) in subpanel A.

To track statistical learning in the brain, we employed neural frequency tagging (***Batterink and Paller, 2017; Choi et al., 2020; Henin et al., 2021***). We quantified the phase coherence of oscillations at the frequency of individual items (present in both Random and Structured blocks) and at half of that frequency reflecting groupings of two items (present only in Structured blocks with category pairs). To measure prediction during encoding, we used multivariate pattern similarity (***Kok et al., 2014, 2017; Demarchi et al., 2019; Aitken et al., 2020***). We first created a template pattern for each scene category based on the neural activity it evoked in visual contacts. We then quantified the expression of these categories during statistical learning, defining prediction as evidence for the second category in a pair evoked by items from the first category. In sum, by assessing iEEG signals during the rapid presentation of scenes, we measured the neural representations underlying statistical learning and prediction, and linked these online learning measures to offline memory, revealing how predictive value constrains memory encoding.

## Materials and Methods

### Participants

We tested 10 participants (7 female; age range: 19-69) who had been surgically implanted with intracranial electrodes for seizure monitoring. Decisions on electrode placement were determined solely by the clinical care team to optimize localization of seizure foci. Participants were recruited through the Yale Comprehensive Epilepsy Center. Participants provided informed consent in a manner approved by the Yale University Human Subjects Committee.

A summary of patient demographics, clinical details, and research participation can be found in ***Table 1***. Given electrode coverage and usable data, we retained 9 patients in the behavioral analyses, 8 patients in the neural frequency tagging analyses, and 7 patients in the neural category evidence analyses.

**Table 1.** Patient Information.

| ID | Age | Sex | nElec (vis) | Implant | Data Collected | Notes |
| --- | --- | --- | --- | --- | --- | --- |
| 1 | 19 | F | 203(21) | R G/S/D | 2S, 2R | R2 mem data not usable (D) |
| 2 | 26 | F | 163(59) | L G/S/D | 2S, 2R | – |
| 3 | 43 | F | 172(10) | Bi D | 1S, 2R | – |
| 4 | 61 | F | 136(0) | Bi D | 1S, 1R | neural mem data not usable (T) |
| 5 | 31 | M | 152(31) | L G/S/D | 2S, 2R | R1 encoding data not usable (T) |
| 6 | 69 | F | 92(7) | L D | 2S, 2R | – |
| 7 | 33 | M | 232(22) | Bi D | 1S, 1R | – |
| 8 | 31 | F | 192(20) | Bi D | 2S, 2R | no mem data collected (C) |
| 9 | 56 | F | 192(36) | Bi D | 2S, 2R | R1 encoding data not usable (T) |
| 10 | 53 | M | 148(0) | Bi D | 2S, 2R | – |

Description of patient participation. ID: patient participation number. Age: in years. Sex: M = Male, F = Female. nElec (vis): the total number of electrode contacts, followed by the number of visual electrode contacts. Implant: R = right-sided implant; L = left-sided implant; Bi = bilateral implant; G = grid; S = strip; D = depth. Data collected: the number of runs for each condition collected (S = Structured, R = Random). Notes: which runs (if any) were excluded from given analyses and why. D = patient distraction (e.g., a clinician coming in and disrupting testing); T = trigger issue (i.e., an error with the equipment such that we could not align individual trials to our neural signal); C = computer error (e.g., the computer crashed).

### iEEG recordings

EEG data were recorded on a NATUS NeuroWorks EEG recording system. Data were collected at a sampling rate of 4096 Hz. Signals were referenced to an electrode chosen by the clinical team to minimize noise in the recording. To synchronize EEG signals with the experimental task, a custom-configured DAQ was used to convert signals from the research computer to 8-bit “triggers” that were inserted into a separate digital channel.

### iEEG preprocessing

iEEG preprocessing was carried out in FieldTrip (***Oostenveld et al., 2011***). A notch filter was applied to remove 60-Hz line noise. No re-referencing was applied, except for one patient, whose reference was in visual cortex, resulting in a visual-evoked response in all electrodes; for this patient, we re-referenced the data to a white matter contact in the left anterior cingulate cortex. Data were downsampled to 256 Hz and segmented into trials using the triggers.

### Electrode selection

Patients’ electrode contact locations were identified using their post-operative CT and MRI scans. Reconstructions were completed in BioImage Suite (***Papademetris et al., 2006***) and were subsequently registered to the patient’s pre-operative MRI scan, resulting in contact locations projected into the patient’s pre-operative space. The resulting files were converted from the Bioimagesuite format (.MGRID) into native space coordinates using FieldTrip functions. The coordinates were then used to create a region of interest (ROI) in FSL (***Jenkinson et al., 2012***), with the coordinates of each contact occupying one voxel in the mask (***Figure 2***).

**Figure 2.**
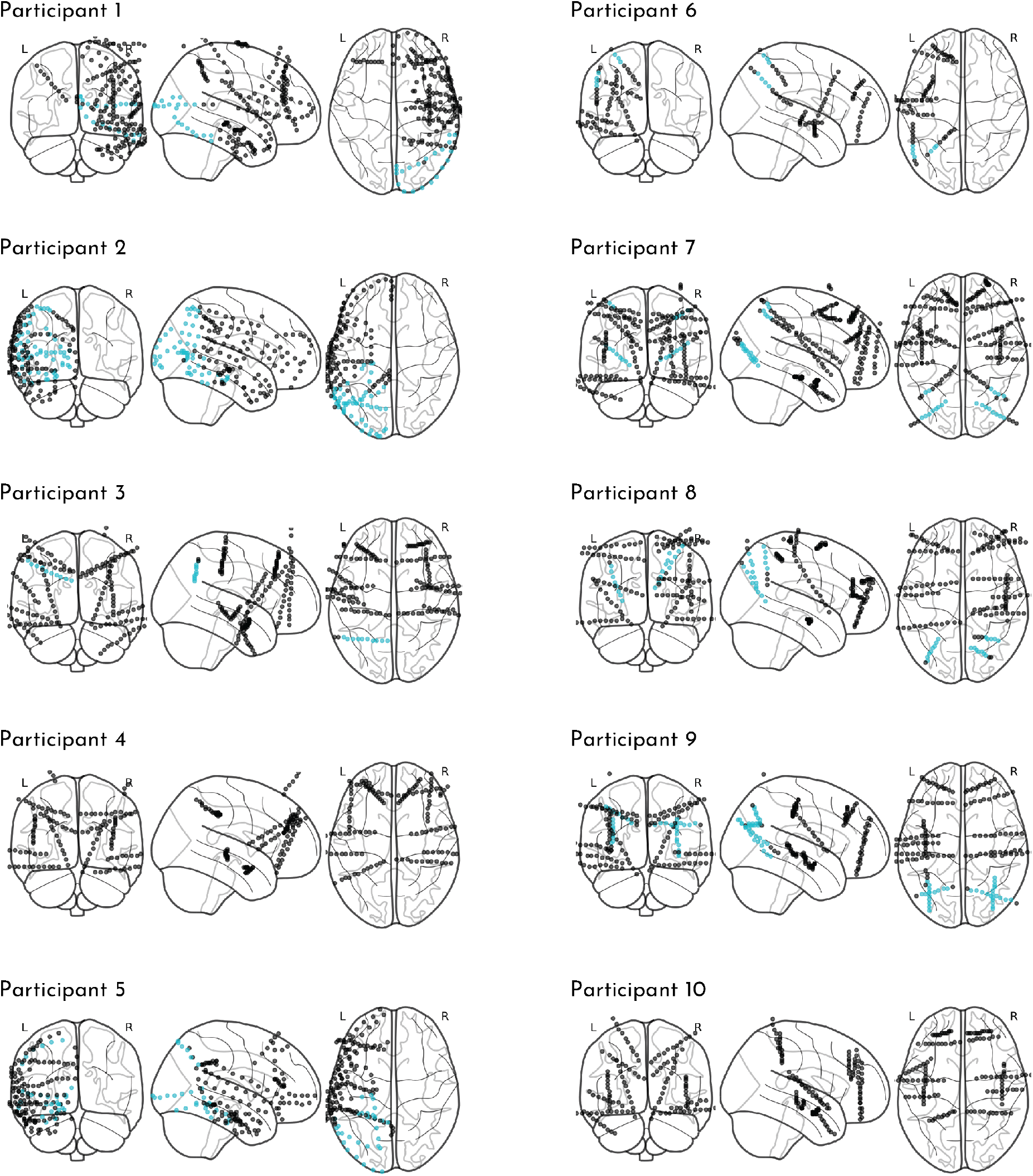
Electrode coverage. The contact locations on the grid, strip, and/or depth electrodes for each participant are plotted as circles in standard brain space. Contacts colored in blue were localized to the visual cortex mask.

For purposes of decoding scene categories, we were specifically interested in examining visually responsive contacts. We defined visual cortex on the MNI T1 2mm standard brain by combining the Occipital Lobe ROI from the MNI Structural Atlas and the following ROIs from the Harvard-Oxford Cortical Structural Atlas: Inferior Temporal Gyrus (temporoocipital part), Lateral Occipital Cortex (superior division), Lateral Occipital Cortex (inferior division), Intracalcarine Cortex, Cuneal Cortex, Parahippocampal Gyrus (posterior division), Lingual Gyrus, Temporal Occipital Fusiform Cortex, Occipital Fusiform Gyrus, Supracalcarine Cortex, Occipital Pole. Each ROI was thresholded at 10% and then concatenated together to create a single mask of visual cortex.

To identify which contacts to include in analyses on a per-patient basis, this standard space visual cortex mask was transformed into each participant’s native space. We registered each patient’s pre-operative anatomical scan to the MNI T1 2mm standard brain template using linear registration (FSL FLIRT (***Jenkinson and Smith, 2001; Jenkinson et al., 2002***)) with 12 degrees of freedom. This registration was then inverted and used to bring the visual cortex mask into each participant’s native space.

In order to ensure that the visual cortex mask captured the anatomical areas we intended, we manually assessed its overlap between the electrodes and made a few manual adjustments to the electrode definition. For example, due to noise in the registrations between post-operative and pre-operative space, as well as from pre-operative space and standard space, some grid and strip contacts appeared slightly outside of the brain, despite being on the surface of the patient’s brain. Thus, contacts such as these were included as “visual” even if they were slightly outside of the bounds of the mask. Additionally, due to the liberal thresholds designed to capture broad visual regions, some portions of the parahippocampal gyrus area contained the hippocampus. Contacts within mask boundaries but clearly in the hippocampus were excluded.

### Procedure

Participants completed the experiment on a MacBook Pro laptop while seated in their hospital bed. The task consisted of up to four runs: two runs of the Structured block and two runs of the Random block. We aimed to collect all four runs from each patient, but required a minimum of one run per condition for subject inclusion. Given that the order of structured vs. random information can impact learning (***Jungé et al., 2007; Gebhart et al., 2009***), the run order was counterbalanced within and across participants (i.e., some participants received Structured-Random-Random-Structured and others Random-Structured-Structured-Random). Participants completed the runs across 1-3 testing sessions based on the amount of testing time available between clinical care, family visits, and rest times.

Each run consisted of an encoding phase and a memory phase. During the encoding phase, participants viewed a rapid stream of scene images, during which they were asked to passively view the scenes. Participants were told that their memory for the scenes would be tested in order to encourage them to pay close attention. Each scene was presented for 267 ms, followed by a 267 ms inter-stimulus interval (ISI) period during which a fixation cross appeared in the center of the screen. These short presentation times were chosen to optimize the task for the frequency tagging analyses, which involves measuring neural entrainment to stimuli.

Within each run, participants viewed a series of images from a set of six scene categories. There were six categories in the Structured block, and six other categories in the Random block. In the Structured block, the scenes categories were paired, such that images from one scene category (A) were always followed by an image from another scene category (B). Thus, A scenes were *predictive* of the category of the upcoming B scenes, or stated another way, the category of B scenes was *predictable* given the preceding A scenes. No scene pairs were allowed to repeat back-to-back in the sequence. In the Random block, all six scene categories (X) could be preceded or followed by any other scene category, making them neither predictive nor predictable. No individual scene categories were allowed to repeat back-to-back.

In total, participants viewed 16 exemplars from each category within each run. To assist patients with remembering these briefly presented images, each individual exemplar was shown four times within a run. Thus, each run was comprised of 16 “subblocks” during which the same set of six exemplar images was repeated four times (384 trials total). Within each subblock, the order of the pairs/images was randomized, with the constraints described above of no back-to-back repetitions. The individual exemplars changed after each subblock, but the category relations were held constant in the Structured block. Participants were not informed of these category pairings, and thus had to acquire them through exposure.

At the end of each run, participants completed a memory test. Participants were presented with all 96 unique images from the encoding phase, intermixed with 24 novel foils from the same categories (4 foils/category). Participants first had to indicate whether the image was old, meaning it was just presented in that run’s encoding phase, or new, meaning that they had not seen that image at all during the experiment. Following their old/new judgment, participants were asked to indicate their confidence in their response, on a scale of 1 (very unsure) to 4 (very sure). Participants had up to 6 s to make each old/new and confidence judgment.

### Frequency tagging analyses

We conducted a phase coherence analysis to identify electrode contacts that entrained to image and pair frequencies (***Henin et al., 2021***). For both Structured and Random blocks, the raw signals were concatenated across runs (if more than one per block type) and then segmented into subblocks comprising 24 trials with the four repetitions per exemplar. We then converted the raw signals for each subblock into the frequency domain via fast Fourier transform and computed the phase coherence across subblocks for each electrode using the formula *R*^2^ = [Σ^*N*^ *cosϕ*]^2^ + [Σ^*N*^ *sinϕ*]^2^. Notably, by computing phase coherence between subblocks, we collapsed over the contribution of individual exemplars that repeated within subblock. In other words, entrainment in this analysis was driven by phase-locking that generalized across exemplars. Phase coherence was computed separately for each contact in the visual cortex mask, and we then averaged across contacts within participant. We focused on phase coherence at the frequency of image presentation (534 ms/image; 1.87 Hz) and pair presentation (1.07 s/pair; 0.93 Hz).

### Category evidence analyses

We employed a multivariate pattern similarity approach to assess the timecourse of category responses. We identified patterns of multivariate activity associated with each category across contacts, frequencies, and time. These category patterns, or “templates”, were defined during the memory phase of the dataset. This was important because the order of categories was random during the memory phase, allowing for an independent assessment of each category across condition regardless of any pairings. We then used these templates to examine category-specific evoked responses during the encoding phase, to assess the presence and timing of category evidence (e.g., for the on-screen category or the upcoming category). The following subsections explain this approach in detail.

#### Frequency decomposition

We employed a Morlet Wavelet approach to decompose raw signals into time-frequency information (***Figure 3A***). We convolved our data with a Complex Morlet Wavelet (cycles = 4) at each of 50 logarithmically spaced frequencies between 2 and 100 Hz to extract the power timecourse at each of these 50 frequencies. This analysis was done separately for each encoding and memory phase of each run, and the data were z-scored across time within each frequency and contact. This procedure was applied across the un-segemented timecourses; we then subsequently carved into trials using the triggers, yielding a vector of frequency and contact information at each timepoint within a trial.

Subsequent analyses required that each trial have the same number of timepoints. However, memory trials were variable lengths, as participants had up to 6 s to respond. There was also slight variability in the encoding trials (most trials were 138 samples long, but some were 136 or 137 samples). To account for this, we considered only the first 138 samples of each memory trial and treated each encoding trial as having 138 samples (interpolating missing timepoints by averaging the last sample of the trial with the first sample of the next trial).

#### Feature selection

We aimed to identify the set of timepoints that produced the best category discrimination. We reasoned that time within a trial would be an important contributor to variance in discriminability, as we would not necessarily expect that timepoints very early on in a trial (immediately after image onset) would produce high discrimination between categories. We also reasoned that the best timepoint(s) may differ from participant to participant depending on their electrode coverage. Therefore, we devised a participant-specific timepoint feature selection process. Importantly, these feature selection steps were conducted within the memory phase data (the same data on which the templates were trained), which were independent of the test data of interest (encoding phase data).

We constructed a set of 30 binary classifiers to distinguish among two categories of a given condition (***Figure 3***B): A1-A2, A1-A3, A1-B1, A1-B2, A1-B3, A2-A3, A2-B1, A2-B2, A2-B3, A3-B1, A3-B2, A3-B3, B1-B2, B1-B3, B2-B3, X1-X2, X1-X3, X1-X4, X1-X5, X1-X6, X2-X3, X2-X4, X2-X5, X2-X6, X3-X4, X3-X5, X3-X6, X4-X5, X4-X6, X5-X6. We employed a linear support vector machine approach using the SVC function in Python’s scikit-learn module, with a penalty parameter of 1.00. We split our data into two-thirds training and one-third test (all within the memory phase), and iterated over the three train-test splits.

**Figure 3.**
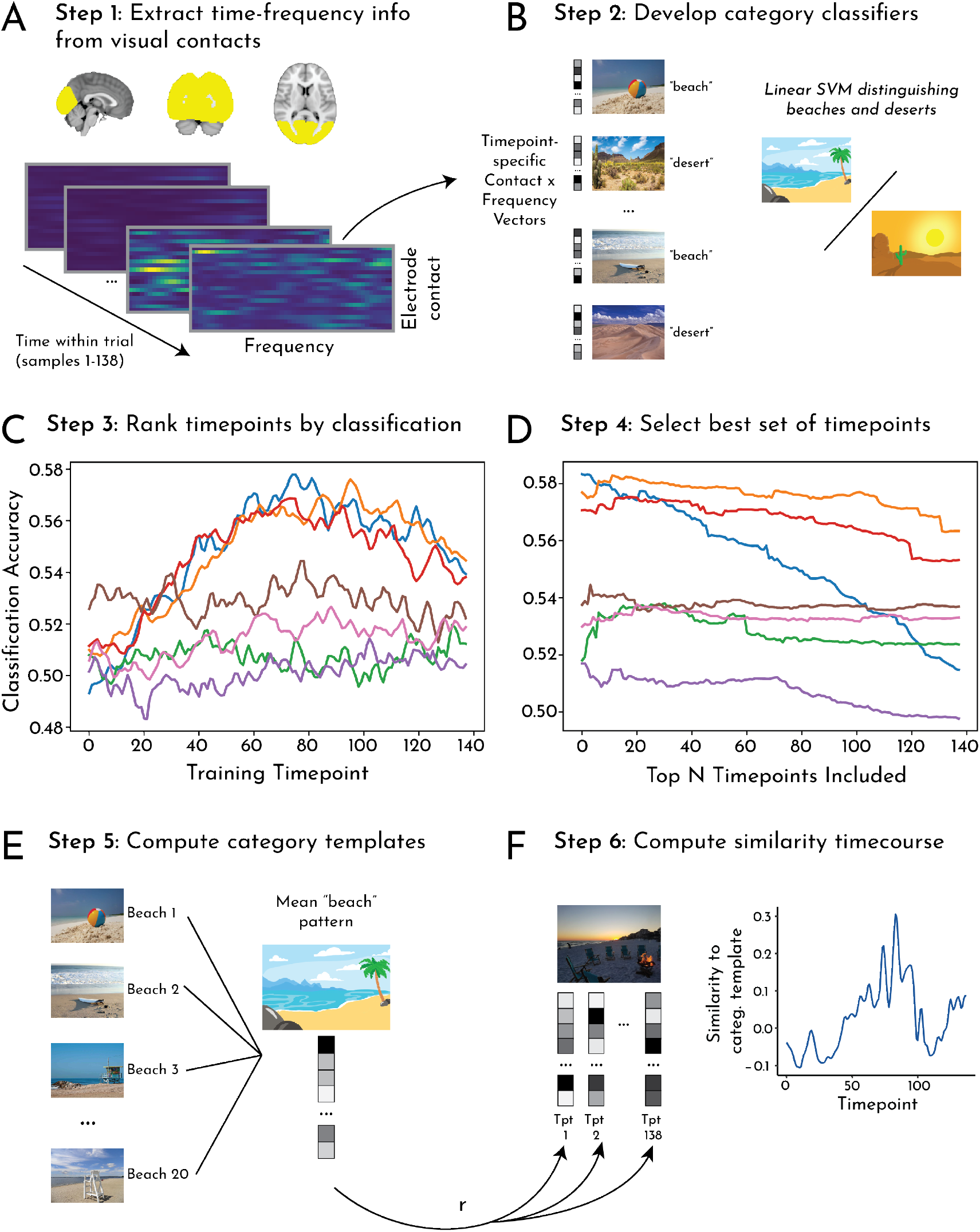
Category evidence analysis pipeline. (A) Step 1: A Morlet wavelet approach was used to extract time-frequency information from contacts in visual cortex. This resulted in contact by frequency vectors for every timepoint of encoding phase and memory phase trials, which served as the neural patterns for subsequent analysis steps. (B) Step 2: To identify the neural patterns that distinguished between categories, we ran a series of binary classifiers for every pair of categories from the memory phase trials. These classifiers were trained on the contact by frequency vectors for a single timepoint (Step 3) or set of timepoints (Step 4). The classifiers were then tested on timepoints from held-out data. (C) Step 3: As a first feature-selection step, we trained and tested the classifiers described in Step 2 separately for all individual timepoints. We then computed the average classification accuracy (across pairwise classifiers) for each timepoint and participant (each colored line indicates one participant). We then ranked the timepoints by classification accuracy. (D) Step 4: To select the *set* of timepoints that produced the best classification for a given participant, we trained and tested the classifiers in Step 2 on an increasing number of timepoints, starting with the best-performing timepoint identified in Step 3 and iteratively adding timepoints by rank. We then computed the per-participant average classification accuracy for each set of timepoints. (E) Step 5: We chose the per-participant top-N timepoint set that produced the best classification accuracy in Step 4, and then averaged contact by frequency vectors across those timepoints (across all exemplars of a given category) to create a “template” of neural activity for each category. (F) Step 6: We then correlated the template for each category from the memory phase with the contact by frequency vector at each timepoint of each trial/exemplar from that category during the (independent) encoding phase, yielding a timecourse of pattern similarity reflecting neural category evidence.

In the first step of feature selection, we independently trained classifiers on a single timepoint (each of the 138 timepoints within a trial) and tested each classifier on all 138 timepoints at test (***Figure 3***C). We averaged the classification over the 138 test timepoints to assess how well training at every timepoint generalized to all other timepoints within a trial. We conducted this analysis for all 30 classifiers and averaged performance over classifiers, yielding a mean classification performance associated with each training timepoint. For each participant, we then computed the rank order of timepoints with respect to their classification, such that the first ranked timepoint was the one that yielded the highest classification, and the last ranked (138th) timepoint is the one that yielded the lowest classification.

To identify the *set* of training timepoints producing the best category classification for a given participant, we repeated the pairwise classification procedure above iteratively training on an increasing number of timepoints, adding from highest to lowest ranked (***Figure 3***D). Thus, these classifiers ranged from training on the single top timepoint, to all 138 timepoints. We again conducted this analysis for all 30 classifiers and averaged performance across them, yielding a mean classification performance associated with the 138 sets of top-N timepoints. We ranked this classification performance again to determine which number of top timepoints produced the highest classification. This number was used to define the templates.

#### Template correlations

Using the set of training timepoints for each participant determined in the feature selection process, we then computed a neural template for each category (***Figure 3***E). We extracted the pattern of activity (i.e., a vector containing electrode contact, time, and frequency) for all instances of a given category during the memory phase, including both old and new images. We then averaged over the timepoints in that participant’s training set. The resulting category pattern vector retained spatial (contact) and frequency information.

To assess the timecourse of neural evidence for a category during the encoding phase, we extracted the pattern of activity (contact and frequency) for each timepoint of every trial of that category (***Figure 3***F). We computed the Pearson correlation between the template and the activity pattern separately for each timepoint within a trial, yielding a timecourse of similarity to the template. The resulting Pearson correlation values were Fisher transformed into *z* values.

We were interested in characterizing the timecourse of a category response not only while that category was on the screen, but also during the surrounding trials. We may observe evidence for a category before it appears, if it can be predicted (as hypothesized for B), or after it disappears, if its representation lingers. Thus, we assessed the timecourse over a window comprising the on-screen category’s trial (“Current”) and the two neighboring trials (“Pre” and “Post” trials). To quantify the response, we subtracted a baseline of average evidence for the other categories of the same condition (e.g., for category A1, how much evidence is there for A1 relative to categories A2 and A3?). For the X categories, which could appear in any order, we ensured that the categories included in the baseline did not appear during the “Pre” and “Post” trials.

We quantified how template similarity changed over time within trial by splitting the trials into “ON” and “ISI” epochs. The ON epoch refers to the part of the trial when the image was on the screen (the first 69 samples, or 267 ms). The ISI epoch refers to the part of the trial after the image disappeared from the screen during the inter-stimulus fixation cross (the second 69 samples, or latter 267 ms).

#### Subsequent memory

To assess how variance in category evidence across trials related to memory outcomes for those trials, we examined predictive and on-screen representations separately for subsequently remembered versus forgotten trials. We conducted this analysis separately for memory of A (as a function of Perceived evidence for A during A and Predicted evidence for B during A) and for memory of B (as a function of Perceived evidence for B during B and Predicted evidence for B during A). Because each image was shown four times, we first averaged the Perceived and Predicted evidence over these four trials. We considered the ISI epoch of each trial, as this was the epoch in which we observed reliable evidence for the Predicted category B during A. As a control analysis, we repeated these steps for the X trials from the Random blocks.

### Statistical analysis

For all analyses (both behavioral and neural), statistical significance was assessed using a random-effects bootstrap resampling approach (***Efron and Tibshirani, 1986***). For each of 10,000 iterations, we randomly resampled participants with replacement and recomputed the mean across participants, to populate a sampling distribution of the effect. This sampling distribution was used to obtain 95% confidence intervals and perform null hypothesis testing. We calculated the *p*-value as the proportion of iterations in which the resampled mean was in the wrong direction (opposite sign) of the true mean; we then multiplied these values by 2 to obtain a two-tailed *p*-value. All resampling was done in R (version 3.4.1), and the random number seed was set to 12345 before each resampling test. This approach is designed to assess the reliability of effects across patients: a significant effect indicates that which patients were resampled on any given iteration did not affect the result, and thus that the patients were interchangeable and the effect reliable across the sample.

## Results

### Memory behavior

We first assessed overall performance in the recognition memory test to verify that participants were able to encode the images into memory. We computed A^′^, a non-parametric measure of sensitivity, from test judgments for items from both Structured and Random blocks. All participants had an A^′^ above the chance level of 0.5 (mean = 0.68; 95% CI = [0.64, 0.70], *p* <0.001; ***Figure 4***A) indicating reliable memory. This was driven by a higher hit rate (mean = 0.51) than false alarm rate (mean = 0.32; difference 95% CI = [0.14, 0.23], *p* <0.001). The proportions of items that were subsequently remembered (hit rate) or forgotten (1-hit rate, or misses) were roughly matched on average, yielding balanced power for within-subject subsequent memory analyses.

**Figure 4.**
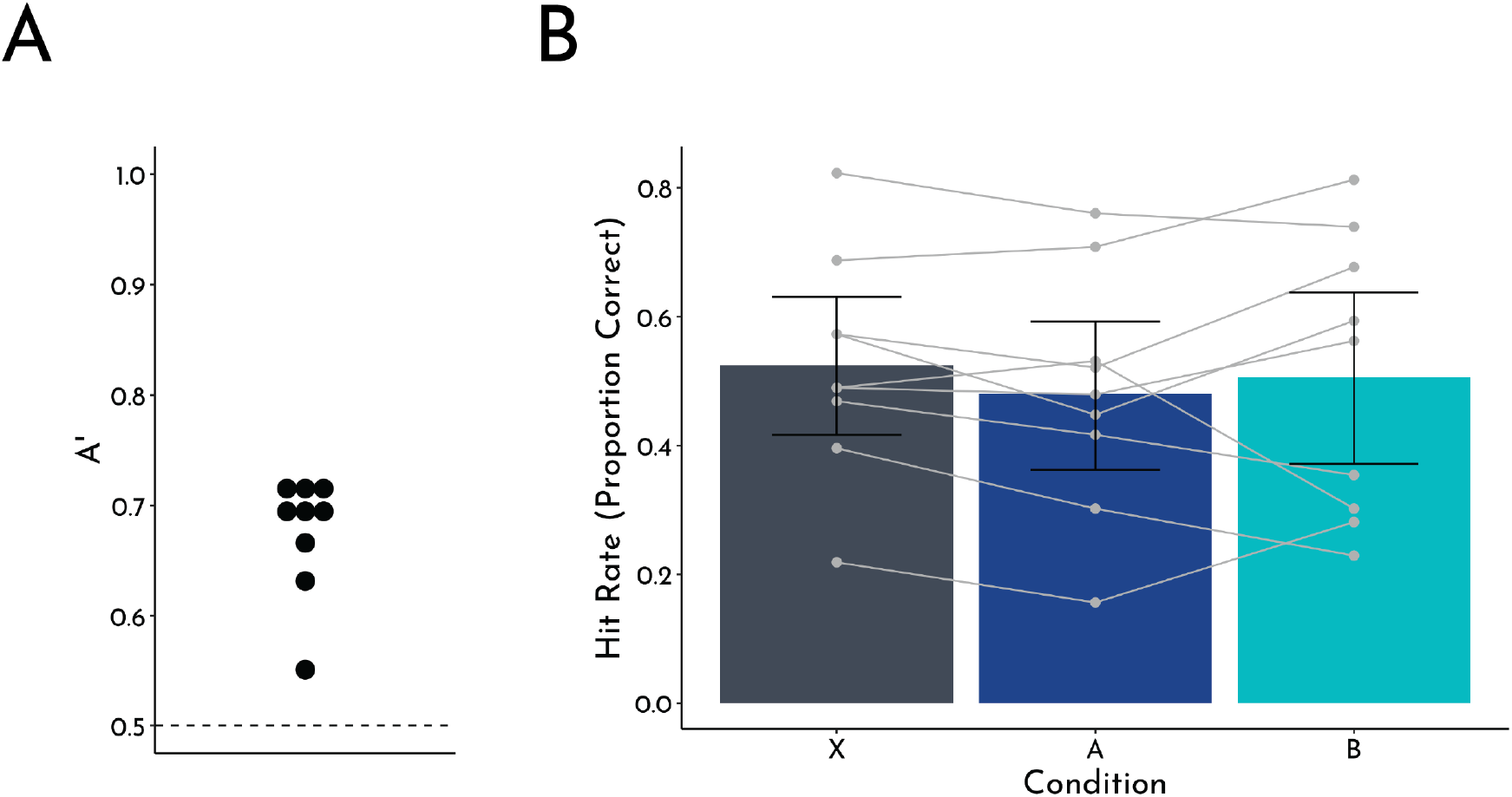
Behavioral results. (A) Overall memory performance collapsed across conditions. A^′^ (a sensitivity measure for recognition memory) is depicted for each participant as a circle. All participants were above chance (0.5). (B) Hit rate as a function of condition (A: predictive; B: predictable; X: control). Group means are plotted as bars, with errors bars representing the bootstrapped 95% confidence interval across participants. Individual participant data are overlaid with the grey circles and lines.

We then assessed how statistical learning affected recognition memory. Based on our prior work (***Sher***man and Turk-Browne, 2020), we hypothesized that the hit rate for items from the predictive A categories in the Structured blocks would be lower than the hit rate for items from the control X categories in the Random blocks. Indeed, we replicated this key behavioral finding (***Figure 4***B), with a significantly lower hit rate for A (mean = 0.48) than X (mean = 0.52; difference 95% CI = [-0.076, -0.010], *p* = 0.012). The hit rate for B (mean = 0.51) did not differ from A (difference 95% CI = [-0.10, 0.059], *p* = 0.51) or X (difference 95% CI = [-0.094, 0.053], *p* = 0.66).

The false alarm rate for X (mean = 0.36) was numerically higher than A (mean = 0.28; difference 95% CI = [-0.0023, 0.16], *p* = 0.064); X was significantly higher than B (mean = 0.29; difference 95% CI = [0.0069, 0.13], *p* = 0.028), though A and B did not differ (difference 95% CI = [-0.074, 0.056], *p* = 0.82). Unlike the higher hit rate for X than A, which was specifically hypothesized based on prior work, the marginally higher false alarm rate for X than A was not expected or consistent with previous experiments. Nevertheless, this complicates interpretation of the hit rate difference as impaired memory for A vs. X. One difference from the prior study is the blocking of Structured (A,B) and Random (X) categories, which may have allowed for differences in strategy or motivation between conditions. Nevertheless, the main memory hypotheses in the current study rest within the A condition (i.e., which A items are remembered vs. forgotten as a function of B prediction), rather than on overall condition-wide differences with X (or B).

### Neural frequency tagging

To provide a neural check of statistical learning of the category pairs in the Structured blocks, we measured entrainment of neural oscillations in visual electrode contacts to the frequency of individual images and image pairs (***Figure 5***A). We expected strong entrainment at the image frequency in both the Structured and Random blocks, as this reflects the periodicity of the sensory stimulation. Critically, we hypothesized that there would be greater entrainment at the pair frequency in Structured compared to Random blocks. This provides a measure of statistical learning because the pairs only exist when participants extract regularities over time in the transition probabilities between categories in the Structured blocks.

**Figure 5.**
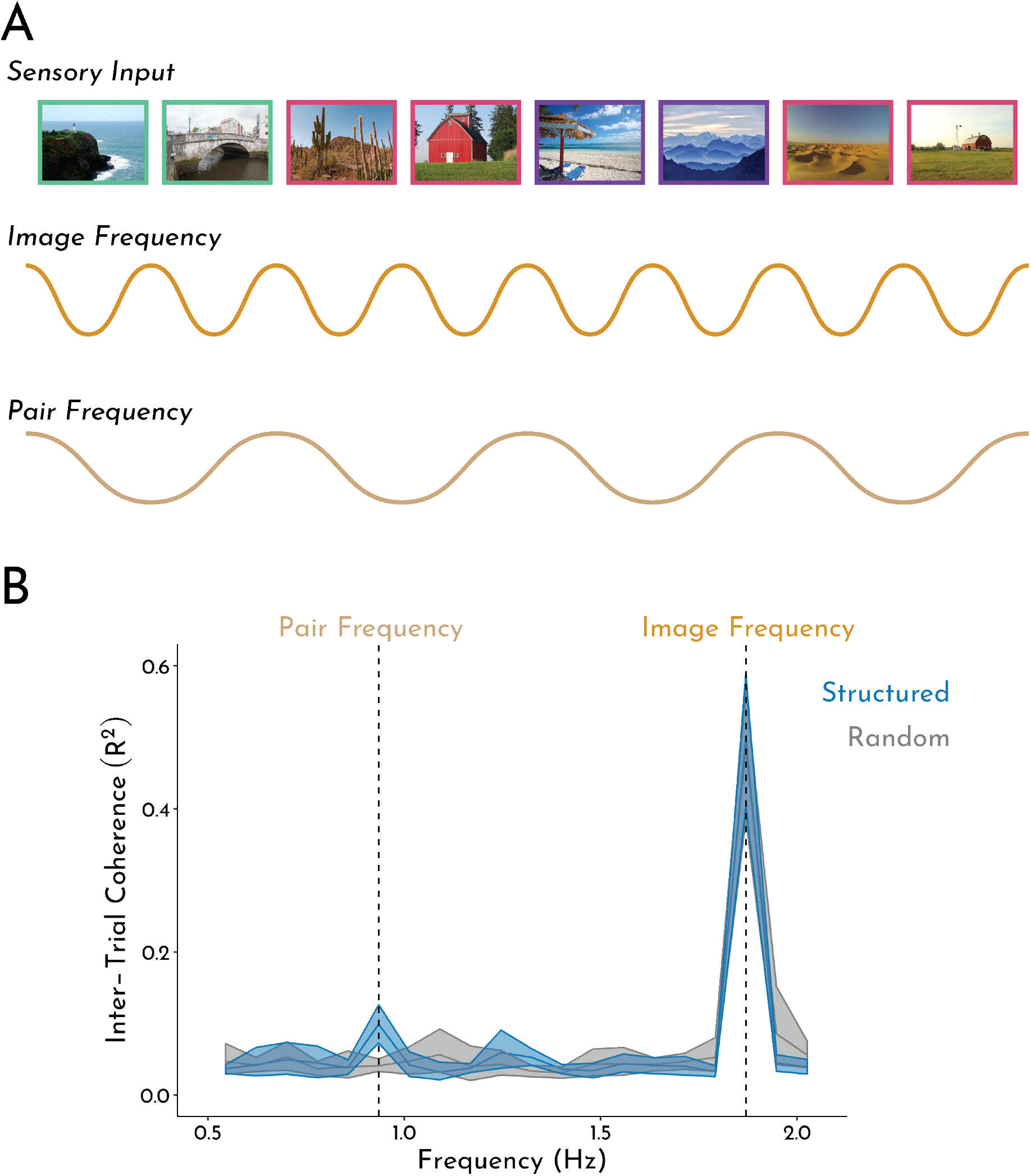
Neural frequency tagging analysis. (A) Schematic of analysis and hypothesized neural oscillations. We expect entrainment of visual contacts at the frequency of images in both blocks. In the Structured block, we also expect entrainment at the frequency of category pairs. (B) These hypotheses were confirmed, with reliable peaks in coherence at the image and pair frequencies in Structured blocks but only at the image frequency in Random blocks. Error bands indicate the 95% bootstrapped confidence intervals across participants.

Consistent with our hypotheses and prior work (***Henin et al., 2021***), there were distinct peaks in phase coherence at both the image and pair frequencies in Structured blocks, but only at the image frequency in Random blocks (***Figure 5***B). To confirm the reliability of these peaks, we contrasted the coherence at the frequency of interest (image: 1.87 Hz; pair: 0.93 Hz) against a baseline of the coherence at frequencies neighboring each of the frequencies of interest (±0.078 Hz). At the image frequency, there were reliable peaks in both the Structured (mean difference = 0.46; 95% CI = [0.37, 0.55], *p* <0.001) and Random blocks (mean difference = 0.42; 95% CI = [0.28, 0.52], *p* <0.001). At the pair frequency, there was a reliable peak in Structured blocks (mean difference = 0.059; 95% CI = [0.035, 0.084]), *p* <0.001), but not Random blocks (mean difference = -0.0027; 95% CI = [-0.016, 0.0085], *p* = 0.68).

Further, the peak in coherence at the pair frequency in Structured blocks was reliably higher than that in Random blocks (mean difference = 0.058; 95% CI = [0.035, 0.083], *p* <0.001), confirming the pair frequency effect was specific to when there was structure in the sequence. There were no differences in coherence at the image frequency across conditions (mean difference = 0.018; 95% CI = [-0.010, 0.048], *p* = 0.25). Together, these results provide strong evidence that visual regions represented the paired categories during statistical learning.

### Neural category evidence

The neural frequency tagging for pairs in Structured blocks indicates statistical learning of the pairs. This learning should create predictive value for the items from the A categories, which afford a prediction of the associated B category. To test for these predictive representations, we employed a multivariate pattern similarity approach that extracted neural evidence for visual categories. For each category, we created a neural template based on the pattern of time-frequency information evoked by each category across visual contacts. We then quantified the expression of these templates in the Structured and Random blocks. As a check, we expected clear neural evidence for the category of the item being presented on the screen.

Critically, we hypothesized that neural evidence for the upcoming B category would manifest before its appearance, in response to an A exemplar. We measured these temporal dynamics of neural category evidence by creating a window of three trials centered on the current item: the trial preceding a trial in which the item appeared (“Pre”), the trial during which the item was on the screen (“Current”), and the trial succeeding the trial in which the item appeared (“Post”). For example, if category Pair 1 involved beaches (A1) being followed by mountains (B1), neural evidence for the mountain category was calculated in response to beach exemplars (Pre), mountain exemplars (Current), and exemplars from the categories that could appear next in the Structured sequence (A2 or A3 categories). These evidence values were averaged across the categories from the same condition (e.g., B1, B2, and B3 for condition B) and plotted over time (***Figure 6***A). For statistical analysis, we averaged the neural category evidence for each category across the timepoints within 6 epochs: when Pre, Current, and Post images were on the screen (“ON”) and during the fixation period between these trials (“ISI”; ***Figure 6***B). We anticipated the evoked response to each image would span ON and ISI periods (as neural processing of the image would take longer than 267 ms), but subdividing in this way allowed us to test for the emergence of predictive evidence of B during the ISI immediately prior to its onset.

**Figure 6.**
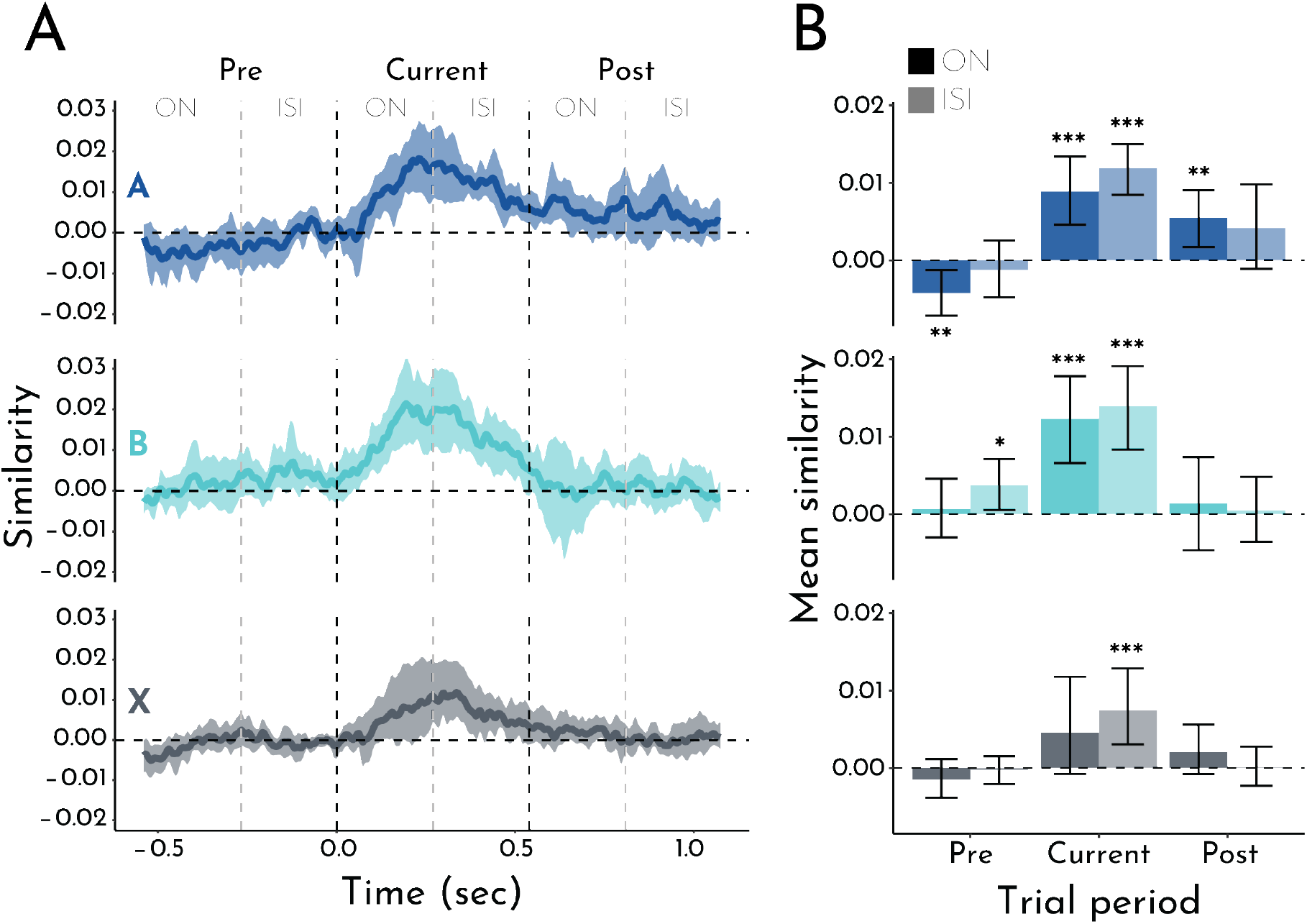
Neural category evidence. (A) Time course of similarity between patterns of neural activity in visual contacts evoked by exemplars from A (predictive), B (predictable), and X (control) categories and category template patterns for A, B, and X. Current refers to the trial when the item was presented, Pre refers to the trial before the item was presented, and Post refers to the trial after the item was presented. For each row/condition, the Pre, Current, and Post trials are compared to the same category template (Current). Error bands reflect the bootstrapped 95% confidence intervals across participants (i.e., any timepoint whose band excludes 0, *p* <0.05). (B) Average pattern similarity collapsed across timepoints within ON (stimulus on screen) and ISI (fixation between stimuli) epochs. Bars represent the means across participants and error bars indicate the bootstrapped 95% confidence intervals. \**p* <0.05; \*\**p* <0.01; \*\*\**p* <0.001

For Current trials (i.e., the trial when the target category was on screen), we found robust (perceptual) evidence for both A and B across both the ON epoch (A: mean = 0.0088; 95% CI = [0.0046, 0.013], *p* <0.001; B: mean = 0.012; 95% CI = [0.0066, 0.018], *p* <0.001) and ISI epoch (A: mean = 0.012; 95% CI = [0.0084, 0.015], *p* <0.001; B: mean = 0.014; 95% CI = [0.0083, 0.019], *p* <0.001). Neural evidence for X categories from Random blocks was not reliable during the ON epoch (mean = 0.0046, 95% CI = [-0.00075, 0.012], *p* = 0.13) but became robust later in the trial during the ISI epoch (mean = 0.0074; 95% CI = [0.0030, 0.013], *p* <0.001). There was greater evidence for B than X categories during both ON (mean difference = 0.0077; 95% CI = [0.00058, 0.015], *p* = 0.031) and ISI epochs (mean difference = 0.0065; 95% CI = [0.00061, 0.012], *p* = 0.031). Considering X as a baseline, this difference shows enhanced perceptual processing of predictable categories. Neural evidence did not differ between A and B categories (*p*s >0.38) or A and X categories (*p*s >0.28).

For Pre trials (i.e., the trial before the target category appeared), we found the hypothesized predictive neural evidence for the B categories during the ISI epoch (just after its paired A category appeared; mean = 0.0037; 95% CI = [0.00054, 0.0071], *p* = 0.019). B evidence was not present during the ON epoch earlier in the Pre trials (while its paired A category was on screen; mean = 0.00063; 95% CI = [-0.0030, 0.0046], *p* = 0.78); this may reflect the time needed for associative reactivation of the B category after perceptual processing of the A item, or anticipation of the timing when B will appear (at the end of the Pre trial). Further supporting our interpretation that Pre evidence of the B categories reflects prediction, no such evidence was observed for X during ON (mean = -0.0015; 95% CI = [-0.0039, 0.0012], *p* = 0.26) or ISI epochs (mean = -0.00031; 95% CI = [-0.0021, 0.0015], *p* = 0.73) or for A during the ISI epoch (mean = -0.0012; 95% CI = [-0.0048, 0.0025], *p* = 0.53). There was *negative* evidence for the upcoming A category during the ON epoch of the Pre trial (mean = -0.0043; 95% CI = [-0.0072, -0.0013], *p* = 0.0052), but this may have been artifactual (see below). When contrasting prediction-related signals across conditions, Pre neural evidence for the B categories during the ISI epoch was reliably greater than X categories (mean difference = 0.0040; 95% CI = [0.00016, 0.0075], *p* = 0.042) and marginally greater than A categories (mean difference = 0.0049; 95% CI = [-0.00051, 0.010], *p* = 0.075).

For Post trials (i.e., the trial after the target category appeared), we found reliable neural evidence for the A categories during the ON epoch (i.e., while its paired B category was on screen; mean = 0.0055; 95% CI = [0.0017, 0.0091], *p* = 0.0018); this effect was not significant during the ISI epoch (mean = 0.0041; 95% CI = [-0.0011, 0.0098], *p* = 0.13). We did not find Post evidence of B or X categories during either ON or ISI epochs (*p*s >0.80), nor was Post evidence for A reliably stronger than B or X (*p*s >0.16). Positive evidence of A during the Post trial may be related to the negative evidence of A during the Pre trial noted above. Because no back-to-back pair repetitions were allowed, in an A1-B1-A2-B2 trial sequence, A1 and A2 were different categories. A1 evidence during B1 was considered a Post trial for the A condition, whereas A2 evidence during B1 was considered a Pre trial for the A condition. Because A1 was one of two baseline categories for A2 (along with the third A category, A3), Post evidence for A1 during B1 would have been subtracted from Pre evidence for A2, leading to a negative effect. We tested this by comparing evidence for A2 (Pre) and A1 (Post) during B1 to the neutral A3 only. This weakened the negative Pre evidence for A, during ON (mean = -0.0027; 95% CI = [-0.0054, 0.00], *p* = 0.058) and ISI epochs (mean = 0.00048; 95% CI = [-0.0022, 0.0038], *p* = 0.82). However, the positive Post evidence for A during the ON epoch remained significant (mean = 0.0081; 95% CI = [0.0036, 0.014], *p* <0.001).

Taken together, these results show that statistical learning of the category pairs in Structured blocks affected neural representations in the task. Not only did visual contacts represent the category of the first and second items in a pair while they were being perceived (A and B evidence during ON and ISI epochs of A and B, respectively), but also the first category during the second (A evidence during ON epoch of B) and the second category during the first (B evidence during ISI epoch after A). This latter effect indicates that the first item in a pair (from A category) had predictive value on average.

### Subsequent memory analysis

We theorized that items with predictive value are a lower priority for new encoding into episodic memory. Here we test this relationship by comparing neural category evidence for remembered vs. forgotten items within participants. That is, although A items had reliable predictive value on average, variability across items may relate to subsequent memory. To the extent that prediction interferes with encoding, we hypothesized that subsequently forgotten A items would elicit evidence for the upcoming B category during their encoding.

Consistent with our hypothesis, B evidence during the ISI epoch after A (i.e., Predicted category) was negatively related to subsequent A memory (***Figure 7***A): forgotten A items yielded reliable B evidence (mean = 0.0092; 95% CI = [0.0023, 0.017], *p* = 0.0030), whereas remembered A items did not (mean = 0.0017; 95% CI = [-0.0016, 0.0049], *p* = 0.31). In contrast, A evidence during the ISI epoch after A (i.e., Perceived category) was reliable for both remembered (mean = 0.012; 95% CI = [0.0091, 0.015], *p* <0.001) and forgotten (mean = 0.014; 95% CI = [0.0077, 0.021], *p* <0.001) A items. This differential effect of subsequent memory on neural evidence for Perceived vs. Predicted categories during the ISI after A was reflected in a significant 2 (evidence category: A, B) by 2 (subsequent memory: remembered, forgotten) interaction (*p* <0.001). This interaction was driven by a marginal difference in neural evidence for the Predicted B category during encoding of subsequently forgotten vs. remembered A items (mean difference = 0.0075; 95% CI = [-0.00046, 0.016], *p* = 0.065), but no reliable difference in neural evidence for the Perceived A category by subsequent memory (mean difference = 0.0022; 95% CI = [-0.0050, 0.0094], *p* = 0.57).

**Figure 7.**
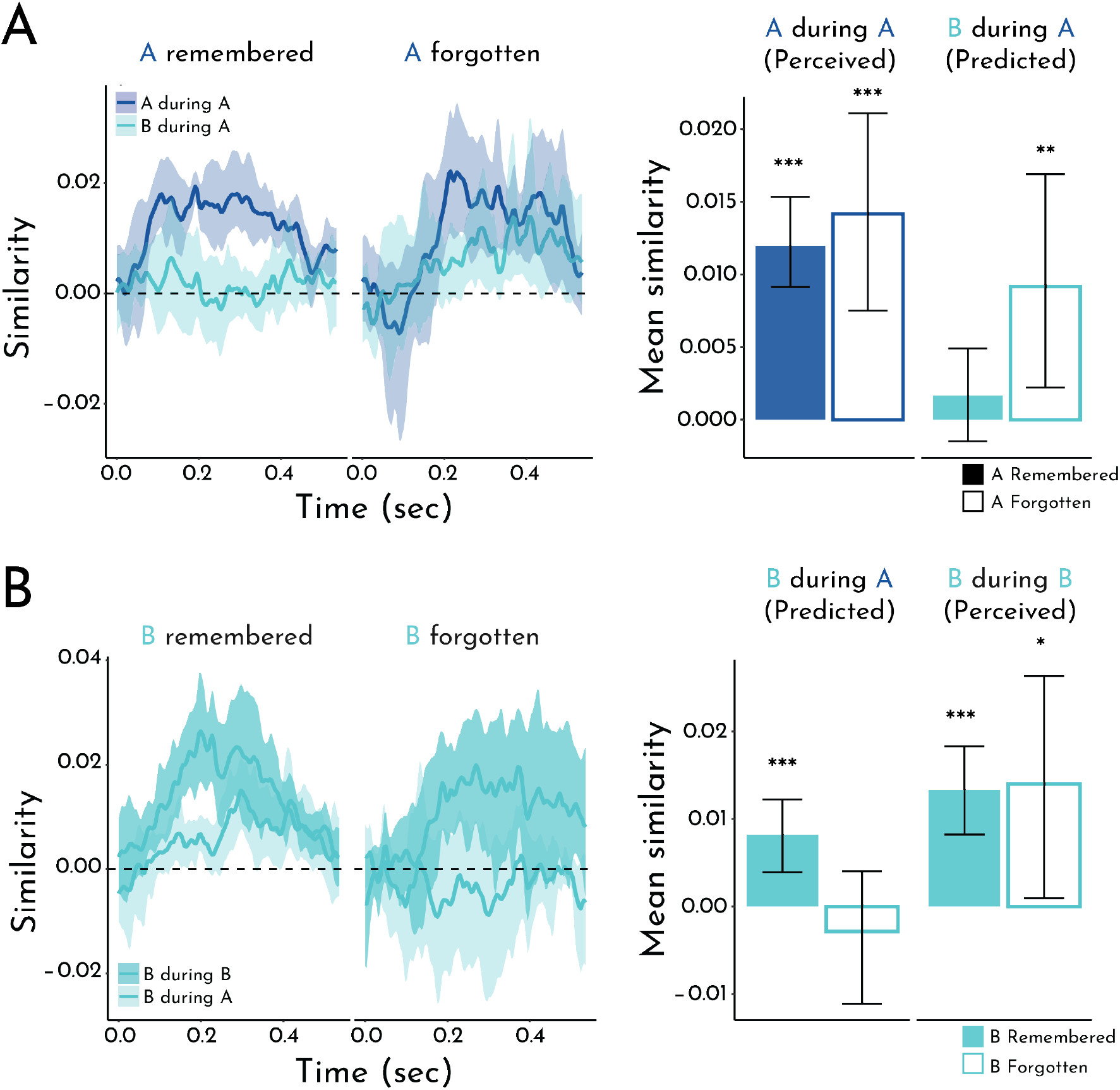
Subsequent memory analysis. A) Left: Timecourse of pattern similarity in visual contacts between A items being encoded and the Perceived A (A during A) and Predicted B (B during A) category templates, as a function of whether A items were subsequently remembered or forgotten. Right: Pattern similarity averaged within the ISI period, the epoch in which we observed overall evidence of prediction, as a function of subsequent memory for A items (filled bars = remembered; empty bars = forgotten). B) Left: Timecourse of pattern similarity in visual contacts between B items being encoded and the Predicted B (B during A) and Perceived B (B during B) category templates, as a function of whether B items were subsequently remembered for forgotten. Right: Pattern similarity averaged within the ISI period, as a function of subsequent memory for B items. Error shading/bars reflect the bootstrapped 95% confidence interval across participants. \**p* <0.05; \*\**p* <0.01; \*\*\**p* <0.001

As a control analysis, we performed the key steps above in the Random blocks. These blocks did not contain pairs, and so we dummy-coded pairs of X items (X_1_-X_2_ instead of A-B). In contrast to Structured blocks, we did not expect that neural evidence of the “Predicted” X_2_ category during the X_1_ ISI would relate to subsequent memory for X_1_. Indeed, there was no reliable evidence for the X_2_ category for either remembered (mean = -0.0029; 95% CI = [-0.0069, 0.00084], *p* = 0.14) or forgotten (mean = 0.0011; 95% CI = [-0.0027, 0.0054], *p* = 0.57) X_1_ items. In contrast, neural evidence for the Perceived X_1_ category during the X_1_ ISI was reliable for both remembered X_1_ items (mean = 0.010; 95% CI = [0.0039, 0.019], *p* <0.001) and forgotten X_1_ items (mean = 0.0065; 95% CI = [0.0022, 0.012], *p* <0.001).

We so far focused on the effects of prediction for memory of the item generating the prediction (A), but what is the mnemonic fate of the item being predicted (B), which in this task with deterministic pairs always appeared as expected? Whereas neural category evidence for B during the A ISI (Predicted) was negatively related to subsequent memory for *A* items, the opposite was true for memory of B items (***Figure 7***B): remembered B items were associated with reliable prediction of B (mean = 0.0082; 95% CI = [0.0036, 0.012], *p* <0.001), but forgotten B items were not (mean = -0.0028; 95% CI = [-0.011, 0.0041], *p* = 0.49). In contrast, and similar to A memory, evidence for B during the B ISI (Perceived) was reliable for both remembered (mean = 0.013; 95% CI = [0.0082, 0.018], *p* <0.001) and forgotten (mean = 0.014; 95% CI = [0.00096, 0.026], *p* = 0.034) B items. We did not find an interaction between category and memory (*p* = 0.22). However, there was a reliable difference in Predicted B evidence for remembered vs. forgotten B items (mean difference = 0.011; 95% CI = [0.00060, 0.021], *p* = 0.039); Perceived B evidence did not differ as a function of memory (mean difference = 0.00064; 95% CI = [-0.014, 0.016], *p* = 0.89).

We repeated the same control analysis of Random blocks, but now focused on subsequent memory for X_2_ items (equivalent to B, rather than X_1_ memory for A). Neural evidence for the “Predicted” X_2_ category during the ISI after X_1_ was not reliable for either remembered (mean = 0.0013; 95% CI = [-0.0020, 0.0043], *p* = 0.44) or forgotten (mean = -0.00048; 95% CI = [-0.0030, 0.0017], *p* = 0.75) X_2_ items.

Together, these results highlight the opposing influence of predictive value on memory for predictive versus predicted items. Namely, prediction of B (during A) is associated with worse memory for predictive A items (suggesting interference between the generation of a prediction and encoding of the current item) but better memory for predicted B items (suggesting that this prediction may potentiate encoding of an upcoming item).

## Discussion

This study demonstrates a trade-off between how well an item is encoded into episodic memory and how strong of a future prediction it generates based on statistical learning. We first used frequency tagging to provide neural verification of statistical learning. During a sequence of scene photographs, electrodes in visual cortex represented pairs of scene categories that reliably followed each other, synchronizing not only to the individual scenes but also to the boundaries between pairs. Next, we used multivariate pattern analysis to assess how the paired categories were represented over time. Items from the first category in a pair elicited a representation of the second category, which grew in strength in advance of the onset of items from the second category. We refer to the ability of an item to generate this predictive representation as its “predictive value”. Critically, by relating these representational dynamics to subsequent memory behavior, we found that forgotten items from the first category triggered reliable predictions during encoding whereas remembered first items had not.

Our work builds upon suggestive evidence from a prior study that predictive value may influence subsequent memory (***Sherman and Turk-Browne, 2020***). This prior study included behavioral and fMRI experiments, whereas the current study employed iEEG. Neural measures are an important advance over behavior alone because they can assay predictive representations during passive viewing at encoding. iEEG is superior to fMRI for this purpose because neural activity is sampled at much greater temporal resolution and activity reflects instantaneous electrical potentials rather than hemodynamic responses smoothed and delayed in time. This provides much greater confidence that the upcoming category was being represented *prior* to its appearance and thus was truly predictive. Moreover, the prior study showed a negative relationship between prediction and memory across participants, whereas the current study established this relationship within participant. This is also an important advance because an across-participant relationship does not provide strong evidence for the claim that prediction during encoding impairs memory. Such a relationship could reflect generic individual differences such that, for example, a participant with better overall memory generates the same weak prediction on both remembered and forgotten trials. In contrast, in this study we were able to link prediction to successful vs. unsuccessful memory formation across items. This more sensitive approach yielded other findings not observed in the prior study, including that memory for B items had an opposite, positive relationship with prediction of B. Taken together, these results provide mechanistic insight into the interaction between predictive value and memory, and speak to theoretical questions about the representations underlying statistical learning and episodic memory.

### Nature of representational changes

Several fMRI studies have shown that statistical and related forms of learning can change neural representations of associated items throughout the human brain (***Schapiro et al., 2012, 2013; Schlichting et al., 2015; Deuker et al., 2016; Tompary and Davachi, 2017***). For example, if exposed to sequential pairs embedded in a continuous stream of objects (akin to the category pairs in the current study), the two objects in a pair come to elicit more similar patterns of fMRI activity from before to after learning, when presented on their own, in the medial temporal lobe cortex and hippocampus (***Schapiro et al., 2012***). Such integration could be interpreted as evidence that the representations of the paired items merged into a single “unitized” representation of the pair that can be evoked by either item (***Fujimichi et al., 2010***). Alternatively, the paired items may remain distinct but become associated, such that either can be reactivated by the other through spreading activation (***Schapiro et al., 2017***). A key difference between these accounts is the timing of how learned representations emerge when one of the items is presented: the merging account predicts that the (same) unitized representation is evoked immediately by either paired item, whereas the associative account predicts that the presented item is represented immediately while the paired item is represented gradually over time through reactivation. These dynamics cannot be distinguished by fMRI because of its slow temporal resolution, but our iEEG approach may shed light.

On the surface, the results of our frequency tagging analysis may seem to suggest a merged representation of the category pairs. The reliable peak in coherence at the frequency of two consecutive stimuli may suggest that electrodes in visual cortex represented the paired categories as a single unit (***Batterink and Paller, 2017***). However, the results of our pattern similarity analysis are more consistent with an association between the paired categories. Although we found that both categories in a pair could be represented at the same time (i.e., predictive B evidence during the A Pre trial and lingering A evidence during the B Post trial, relative to no such evidence on X trials), these representations were offset in time. The representation of the A category was robust during both the ON and ISI epochs of the A trial, whereas the representation of the B category was not reliable during the ON epoch and only emerged during the ISI epoch. Thus, our results are more consistent with an associative account in visual cortex. It remains possible that the hippocampus or other brain structures represent statistical regularities through unitized representations. Moreover, one limitation of our study is that we did not measure representations of individual categories before and after learning to directly assess representational change. Though note that this is more important for fMRI where, unlike with iEEG, the coarse temporal resolution makes it difficult to separate neural responses of paired stimuli during statistical learning.

### Predictive interference on memory encoding

The timecourse of predictive representations also sheds light on the temporal dynamics of the interaction between episodic memory and statistical learning. When examining the overall effect of prediction, we found reliable B evidence during the ISI epoch of A, immediately preceding the appearance of B. However, this result was obtained by averaging across all trials, both remembered and forgotten. Thus, it was possible that when separated out by subsequent memory, a different pattern would emerge. One possibility is that B evidence would come online earlier for forgotten items, which might suggest that the observed impairment in A memory resulted from interference with perceptual processing of A. To the contrary, the difference in B evidence for remembered vs. forgotten A items was clearest during the ISI after A was removed from the screen, which suggests that prediction may interfere with later, post-perceptual stages of processing to impair encoding.

Interestingly, evidence for the current A category was comparable across remembered and forgotten A items. Thus, in this paradigm, variance in memory was explained solely by prediction of the upcoming category, not the strength of perceptual processing of the category being encoded (***Kuhl et al., 2012***) nor modulation of this processing by prediction (both of which would have affected A evidence). The lack of a relationship between A evidence and A memory may reflect a tradeoff: category evidence may reflect representation of the most diagnostic features of a category, which would enhance memory for these features while impairing memory for idiosyncratic features of particular exemplars. A related account may explain why predictive B evidence was positively linked to B memory (***Smith et al., 2013; Thavabalasingam et al., 2016***): B evidence during the A ISI may potentiate the diagnostic features of the B category, enhancing the salience of idiosyncratic features of B when it appears to strengthen episodic memory for B. Future studies could test these possibilities by using a more continuous measure of memory precision and by testing on modified items that retain category-diagnostic vs. idiosyncratic features.

This work builds on existing theories considering the complex interplay between memory encoding and memory retrieval. To the extent that prediction from statistical learning can be considered as “retrieval” of an associated memory (***Kok and Turk-Browne, 2018; Hindy et al., 2016***), our findings converge with the notion that the brain cycles between mutually exclusive encoding and retrieval states (***Hasselmo et al., 2002; Duncan et al., 2012; Long and Kuhl, 2019; Bein et al., 2020***). Further, a recent computational model suggests that predictive uncertainty determines when memories should be encoded or retrieved (***Lu et al., 2022***). The model accounts for findings that familiar experiences are more likely to evoke retrieval (***Patil and Duncan, 2018***), and thus may help to explain why predictions from statistical learning are prioritized over episodic encoding.

### Neural source of predictions

The current study sought to decode evidence of visual categories and so focused on electrode contacts in visual cortex. This adds to a growing literature on predictive signals in visual cortex (***De Lange et al., 2018; Kim et al., 2020***). However, these signals may originate elsewhere in the brain. A strong candidate is the hippocampus and surrounding medial temporal lobe cortex. In addition to representing predictions (***Kok and Turk-Browne, 2018; Sherman and Turk-Browne, 2020***), the hippocampus interfaces between perception and memory (***Treder et al., 2021***) and has been shown to drive reinstatement of predicted information in visual cortex (***Bosch et al., 2014; Tanaka et al., 2014; Danker et al., 2017; Hindy et al., 2016***).

Beyond generating predictions, the hippocampus may also be the nexus of the interaction between episodic memory and statistical learning, given its fundamental role in both functions (***Schapiro et al., 2017***). Indeed, given the necessity of the hippocampus for episodic memory, our study raises questions about how the representations of perceived and predicted categories in visual cortex are routed into the hippocampus for encoding. One intriguing possibility is that these representations are prioritized according to biased competition (***Desimone, 1998; Hutchinson et al., 2016***), leading to preferential routing and subsequent encoding of predicted, but not perceived, information in the hippocampus. Relatedly, recent work had found that encoding vs. retrieval states are associated with distinct patterns of activity in visual cortex (***Long and Kuhl, 2021***), suggesting that representations in visual regions may be fundamentally shaped by memory state in the hippocampus.

The patients in the current study had relatively few contacts in the hippocampus and medial temporal lobe cortex, precluding careful analysis of prediction in these regions and how it relates to visual cortex. Future studies with a larger cohort of patients and/or high-density hippocampal recordings would be useful for this purpose. Likewise, future studies could disrupt the hippocampus through stimulation to establish its causal role in predictive representations in visual cortex.

## Conclusion

In examining the trade-off between prediction and memory encoding, our work suggests a novel theoretical perspective on why predictive value shapes memory. We argue that because memory is capacity- and resource-limited, memory systems must prioritize which information to encode. When prior statistical learning enables useful prediction of an upcoming experience, that prediction takes precedence over encoding. In this way, encoding is focused adaptively on experiences for which there is room to develop stronger predictions.

## Acknowledgments

We are grateful to the patients who participated in this study. We thank Kun Wu for providing the electrode reconstructions, Christopher Benjamin for helping to recruit patients and coordinate testing, Richard Aslin and Sami Yousif for helpful conversations, and Gregory McCarthy for advice about data collection and analysis, as well as for feedback on the manuscript. This work was supported by NIH grant R01 MH069456 (N.B.T-B.), the Canadian Institute for Advanced Research (N.B.T-B.), and an NSF GRFP grant (B.E.S).

## Competing Interests

The authors declare no competing interests.

## Notes

### Competing Interest Statement

The authors have declared no competing interest.

